# Does sex influence element accumulation in honey bees: workers vs drones

**DOI:** 10.1101/2023.08.28.555195

**Authors:** Nenad M. Zarić, Robert Brodschneider, Walter Goessler

**Affiliations:** University of Belgrade - Faculty of Biology, Studentski trg 16, 11000 Belgrade, Serbia; University of Graz, Institute of Chemistry, Analytical Chemistry for Health and Environment, Universitaetsplatz 1, 8010 Graz, Austria; University of Graz, Institute of Biology, Universitaetsplatz 2, Graz, 8010, Austria

**Keywords:** Apis mellifera, element composition, food filtration, ICPMS

## Abstract

Honey bees are social insects that have divided their work. Bees have two sexes, male and female. Female honey bees are queens and worker bees, while males are called drones. Worker bees have different tasks in the hive including gathering of food, its processing, caring for brood, protecting the hive, produce wax, nourish workers, drones and queen. Drones are male bees whose only role is to mate with a virgin queen. Many studies have dealt with physiology, behavior, morphology of workers and drones. This is the first study that compared differences in element accumulation and composition between workers and drones honey bees. It was observed that worker honey bees have higher concentration, from 116 % to 517 %, of most elements analyzed. Drones had higher concentration of essential elements to bees, Na, Mg, P, S, K, Zn, Cu and especially Se which was 2.2-fold higher. These differences can be attributed to environmental exposure; reproductive role of drones, high Se in sperm; but mostly to the food workers and drones eat. Worker bees feed on unprocessed food, mostly pollen, which is rich in minerals. Drones are fed by worker bees pre-processed food. Here we show that worker honey bees have a filtering mechanism that enables them to accumulate non-essential elements and not pass them on through the processed food to drones or larva.

## Introduction

Origin of metals in the environment can be natural or anthropogenic. They are non-degradable, meaning that they can only change their chemical form and enter biological systems (Perugini et al., 2011). Essential elements that have a known metabolic function are Na, K, Ca, Mg, and P (Nation, 2015). In addition, some metals are essential parts as enzyme co-factors. These include Cu, Mo, Co and Cr (Gordon, 1959). Zn and Mn are essential for hardening of mandibles cuticle (Behmer, 2008). A central element in cytochrome enzymes is Fe (Behmer, 2008). All of these elements are considered essential, however their exact roles and amounts needed in insects and especially honey bees have not been studied in detail. For some of the elements, for example Se, it is known that lower doses can be very beneficial, but higher doses are toxic (Alburaki et al., 2019). Some elements, such as Al, Ba, Cd, Ni, Pb and Sr are considered non-essential, and might interact with macromolecules by replacing essential metals and could be toxic.

The main reason for managing honey bee colonies is their pollination services. With the growth of population so will grow the dement for food, and hence pollination services. In the last few decades there is a trend in decline of honey bees and this has been named colony collapse disorder (CCD). There are a number of factors that influence this: habitat fragmentation, pathogens, climate change, intensive agriculture, pollution (Potts et al., 2010; Vanbergen & Initiative, 2013). This includes the use of pesticides and emission of toxic metals.

Honey bees are social insects that are generally divided into three groups. The queen, usually only one in the hive, is a reproductive female. Its job is to lay eggs and regulate the hive’s activity, throughout emission of pheromones (Pankiw et al., 1998). The queens are developed from larvae selected by worker bees that are specially fed an exclusive food called “royal jelly”. Queens are largest members of the bee hive. The second group are workers, non-reproductive females. They are developed from fertilized eggs, but are not sexually developed. Workers, depending on their age, have different roles in the hive. They take care of the brood, build and protect the hive, forage for food and water, process food, nourish the workers, drones and queens, produce was, etc. (Schmickl & Crailsheim, 2002, 2004). To be able to perform all these tasks workers are equipped with well-developed hypopharyngeal and mandibular glands, wax glands and scent glands (Hrassnigg & Crailsheim, 2005). The third group are male bees called drones, grown from unfertilized eggs. Usually there are a few hundred drones in the hive during spring and summer. Their only role is to produce sperm and mate with a virgin queen from a different hive then the one they came from.

Metals and metalloids in honey bees have been the subject of many studies. Most of these studies used honey bees to monitor metal and metalloid pollution in the environment (Fry et al., 2023; Hladun et al., 2016; van der Steen et al., 2016; Zarić et al., 2018; Zaric et al., 2018; Zarić et al., 2022; Zhou et al., 2018). Some focused on essential and non-essential elements, not just in whole honey bees, but in their hemolymph as well (Ilijević et al., 2021). However, there is only one previous study done on elemental composition of individual bees (Zarić et al., 2021).

Differences in physiology, behavior, morphology and nutrition between workers and drones of honey bees has been studied before (Brodschneider & Crailsheim, 2010; Henderson, 1992; Hrassnigg & Crailsheim, 2005). However, the differences in element accumulation and composition were not studied in any adult insects until now. The aim of this study is to determine the differences in element composition between male (drone) and female (worker) honey bees. For the purpose of this study individual foragers and drones were analyzed for their content of 31 different elements. This will help explain how different lifecycles of females and males influence their element composition.

## Materials and Methods

### Sample collection

Samples honey bees were taken from a hive located in Gries, Graz, Austria. The apiary was located in the city center on a building roof. Samples of adult worker honey bees were collected from the outer most frame that had honey on it but no brood, as it is believed that these are forager bees that have already flown out of the hive (Medrzycki et al., 2013). The foragers and drones were collected from the same hive. All of the samples were collected on the same day in June 2021. Individual bees (forager or worker) were placed into separate Eppendorf 2 mL tubes. After collection they were frozen at -80°C and kept in the freezer until analyses.

### Sample preparation

The samples were freeze-dried before analyses. Afterwards individual forager or drone honey bees were weight into clean 10 mL quartz vessels. The digestion was done using an ultraCLAVE IV microwave digestion system (MLS GmbH, Leutkirch, Germany) with 2 mL conc. HNO_3_ and 3 mL ultrapure water (Supplementary material). With each digestion three digestion blanks (2 mL conc. HNO_3_ and 3 mL ultrapure water) and three reference materials 8414 “Bovine muscle powder” (NRC, Canada) (∼0.25 g and 5 mL conc. HNO3) were analyzed. After the loading pressure of 40 bars inside the vessels was achieved by high purity Argon 5.0, the microwave heating program was started: The temperature was raised gradually to 80 °C in 10 min, ramped to 150 °C in further 25 min, then ramped to 250 °C in 20 min and finally held at 250 °C for 30 min. After cooling the digestion solutions were transferred to 50 mL Cellstar tubes and diluted with ultrapure water to a final volume for blanks and samples of 20 mL and reference material 50 mL (10% (*v/v*) nitric acid).

### Determination of element concentrations

All element concentrations were determined using inductively coupled plasma mass spectrometry - ICPMS (Agilent ICPMS 7700x, Waldbronn, Germany). For twenty-seven elements an external calibration curve in four different concentration ranges and with six points each, was made in 10% HNO_3_ (0.0100–5.00 μg L^−1^ for Li, V, Cr, Co, Ni, As, Se, Mo, Ag, Cd, Sn, Sb, Cs, Tl, Pb and U; 0.1–50 μg L− 1 for B, Ba, Cu, Rb, and Sr; 1.00–500 μg L^−1^ for Al, Mn, Fe, Zn; 100–50,000 μg L^−1^ for Na, K, Ca, Mg, P and S). Instrument performance is reported in Table S1. Selected mass, tune mode, internal standard for correction for each element analyzed are reported in Table S2.

### Quality control

Internal standard solution containing 200 μg L^−1^ of Be, Ge, In and Lu in a matrix of 1% v/v HNO_3_ was continuously added for instrument stability control. In addition, drift standards were measured every ten samples. The accuracy was evaluated using two reference materials: SRM 1640a Trace elements in natural water (National Institute of Standards & Technology, Gaithersburg, USA) and CRM 8414 Bovine muscle powder (NRC, Canada) (Supplementary material, Table S3 and S4).

### Statistical analyses

For statistical analysis Microsoft Excel 2021 and IBM SPSS Statistics 27 were used. To assess statistically significant differences between female (foragers) and male (drones) honey bees both parametric – Independent t-test, and non-parametric – Kruskal-Wallis H followed by Dunn-Bonferroni post hoc test were applied to the dataset.

## Results and discussion

Average dry weight of collected worker honey bees was 49 ± 14 mg, which is higher than in our previous study (29.2 ± 5.8 mg) for bees from Serbia (Zarić et al., 2021), or the one done by (Brodschneider et al., 2009)) in Graz. Drones average dry weight was 63 ± 4 mg, which is a bit higher than reported in the literature to vary from 30.7 mg to 56.9 mg (Henderson, 1992). In this study drones have approximately 30% higher dry weight compared to worker bees. There were no literature data for dry weight comparison, however fresh weight drones are usually twice heavier compared to worker bees (Es’kov & Es’kova, 2013).

Out of 32 analyzed elements 27 were above the detection limit (LOD). Elements below LOD were discarded from further discussion. The three most abundant elements in both workers and drones are K > P > S, while the lowest concentrations were observed for Sb > Ag (Table 1). Most of the analyzed elements had statistically significant differences between workers and drones in both parametric and non-parametric tests (Supplementary material, Table S5). The only elements that did not show statistically significant differences were B, Cr and Sn, although all of them had higher concentrations in workers.

**Table 1.** Concentration of elements (mg kg^-1^ dry weight ± standard deviation (SD)) in worker (n = 17) and drone (n = 12) honey bees.

| Element | Worker | Drone |
| --- | --- | --- |
| Ag** | $0.020 \pm 0.011$ | $0.0057 \pm 0.0016$ |
| Al** | $17.4 \pm 7.6$ | $10.2 \pm 4.7$ |
| As** | $0.0700 \pm 0.029$ | $0.037 \pm 0.017$ |
| B | $4.6 \pm 1.2$ | $4.4 \pm 1.4$ |
| Ba** | $1.55 \pm 0.57$ | $0.68 \pm 0.32$ |
| Ca** | $872 \pm 280$ | $582 \pm 119$ |
| Cd** | $0.062 \pm 0.035$ | $0.0126 \pm 0.0079$ |
| Co** | $0.066 \pm 0.028$ | $0.032 \pm 0.015$ |
| Cr | $0.16 \pm 0.13$ | $0.099 \pm 0.062$ |
| Cu* | $21.6 \pm 6.4$ | $26.1 \pm 2.8$ |
| Fe** | $145 \pm 47$ | $95 \pm 21$ |
| K* | $8310 \pm 1509$ | $9557 \pm 1587$ |
| Mg* | $755 \pm 192$ | $892 \pm 81$ |
| Mn** | $59 \pm 36$ | $12.2 \pm 5.0$ |
| Mo** | $0.79 \pm 0.25$ | $0.273 \pm 0.048$ |
| Na** | $423 \pm 109$ | $813 \pm 176$ |
| Ni* | $0.30 \pm 0.21$ | $0.149 \pm 0.048$ |
| P** | $5494 \pm 1395$ | $7888 \pm 687$ |
| Pb** | $0.183 \pm 0.076$ | $0.081 \pm 0.035$ |
| Rb* | $9.2 \pm 2.6$ | $6.2 \pm 3.8$ |
| S** | $3168 \pm 832$ | $5183 \pm 378$ |
| Sb** | $0.030 \pm 0.017$ | $0.0107 \pm 0.0038$ |
| Se** | $0.233 \pm 0.052$ | $0.518 \pm 0.071$ |
| Sn | $0.036 \pm 0.018$ | $0.0342 \pm 0.0097$ |
| Sr** | $1.26 \pm 0.41$ | $0.62 \pm 0.32$ |
| V** | $0.042 \pm 0.021$ | $0.0181 \pm 0.0077$ |
| Zn* | $80 \pm 30$ | $107 \pm 18$ |

Independent t-test \* $p < 0.05$ ; \*\* $p < 0.01$

Statistically significant higher concentrations in worker bees compared to drones were observed for Al, Ca, V, Mn, Fe, Co, Ni, As, Rb, Sr, Mo, Ag, Cd, Sb, Ba and Pb (Fig. 1). The worker honey bees that were sampled are mostly foragers. These are the bees that fly out 12-15 times per day to gather food and water for the hive (Perugini et al., 2011). These bees have been exposed to the full impact of the environment and the pollution present in water, soil (through plant pollen and nectar) and air (Hladun et al., 2015; Sadeghi et al., 2012; Zarić et al., 2017).

**Figure 1.**
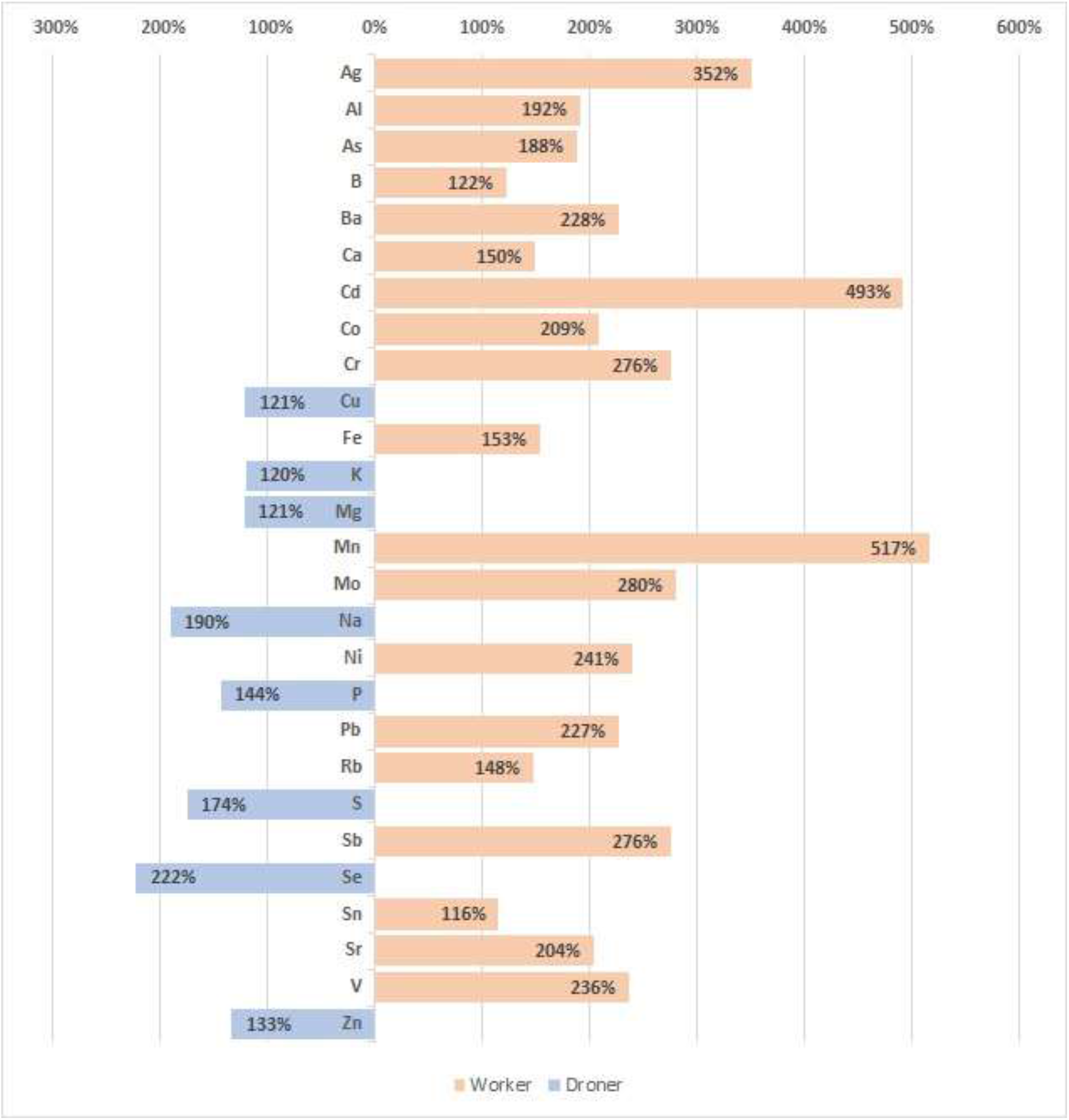
Percent differences in element concentrations between worker and drone bees

Higher concentration in drones compared to foragers can be observed for Na, Mg, P, S, K, Cu, Zn and Se. For insects Na, Mg, P and K are considered essential (Nation, 2015). Cu is an important part of enzymes (Gordon, 1959). Out of all the elements that had higher concentrations in drones, Se was the one with the biggest difference. It was more than 2-fold higher in drones compared to foragers. It was proven that Se is very important for fertility and sperm quality in many animals and man (Alavi et al., 2020; Hansen & Deguchi, 1996). For insects Se could be beneficial for egg fertilization (Martin-Romero et al., 2001). Considering that drones only role is to produce sperm and mate with a queen, we assume that higher Se content is due to its accumulation in their sperm.

Lower concentrations of most elements in drones compared to worker bees is likely due to their lifecycle. Drones fly out during mating season only once per day, and only if the weather conditions are optimal (Hrassnigg & Crailsheim, 2005). If they do not mate within 30 minutes they return to the hive. In comparison to foragers that spend most of the day outside of the hive gathering food, drones are most of the day inside. They are not as much exposed to outside environmental pollution.

In a recent study by Taylor et al. (2023) it was concluded that elements are not attached to the surface of the bee, but are bioaccumulated in the honey bee body. This was confirmed by our own experiments on washed bees. Most of the elements honey bees accumulate are from the food they eat. While worker bees eat unprocessed food, drones are fed nectar or honey and protein jelly. Drones are missing hypopharyngeal glands (glands that produce food), wax glands, and most of the structures to collect food (Hrassnigg & Crailsheim, 2005). They also have a slenderer honey stomach compared to workers. Drones consume only 2-3% of pollen that worker bees do (Szolderits & Crailsheim, 1993). Most of the food that drones get is pre-digested, via proteinaceous glandular secretions and honey provided by workers. Larva are also fed only processed and “filtered” food provided to them by nurser bees. From our own preliminary data, we observed that all the non-essential elements that had lower concentrations in drones, also had very low concentrations in larva. Worker honey bees, especially nurse bees, obviously have a mechanism for filtering unwanted elements from food, in this case pollen. These elements are accumulated in the worker honey bee and are not passed on to the food they produce which is fed to drones and larva. Essential elements are most likely not filtered by worker bees, and thus not accumulated in them, and that is the reason their concentrations are higher in drones and larva. The mechanism enabling worker bees to filter non-essential elements from their food should be studied further.

## Conclusions

This work shows that there are differences in element accumulation between sexes of honey bees. Statistically significand differences were observed for 24 out of 27 detected elements. Drones had higher concentration only for essential elements, Na, Mg, P, S, K, Zn, Cu and Se. The rest of the elements had significantly higher concentrations in worker bees. Se is known to be important for sperm quality and fertility in many animals and humans. This is the first time it was observed that male insects have higher Se content compared to females. For the rest of the elements, a couple of factors could influence these differences. Sampled bees were mostly foragers that spend most of their time outside of the hive gathering food. Hence, they are more exposed to environmental pollution, compared to drones that spend most of their life inside the hive. However, most likely explanation is in the food they consume. Worker honey bees feed on unprocessed food from the environment, mostly pollen, which is rich in minerals. Drones on the other hand are fed pre-digested, “filtered” food produced by worker bees. The underlying mechanism of filtering noon-essential elements in honey bees is still unknown and needs farther study.

## Acknowledgments

This paper was made possible through a bilateral project between Serbia and Austria No. 337-00-577/2021-09/19 and No. RS 17/2022. The authors acknowledge the financial support by the University of Graz. We also acknowledge the financial support of the Ministry of Science, Technological Development and Innovation of the Republic of Serbia (contract No. 451-03-47/2023-01/ 200178).

