## Supplementary material for "Does sex influence element accumulation in honey bees: workers vs drones"

Text S1:

Chemicals and standards

Purification system (Milli-Q, Merck Millipore, Darmstadt, Germany) was used to provide purified water (18.2 MΩ cm). Nitric acid ($HNO_3$) Rotipuran p. a. ≥ 65% (Carl Roth, Karlsruhe, Germany) was subboiled with a MLS duoPUR (MLS, Leutkirch, Germany) prior to its use for the preparation of samples. For internal standards and preparation of calibration standards we used ICP Single-Element Standards Certipur (Merck Millipore, Darmstadt, Germany) and Single Element Standards for ICP (Carl Roth, Karlsruhe, Germany). Fifteen and fifty mL Cellstar polypropylene tubes (Greiner Bio-One International GmbH, Kremsmünster, Austria) were used for preparation of all solutions.

Table S1. Performance of the ICPMS

| **Parameter** | **No-gas mode** | **Collision mode** | **Reaction mode** |
|---|---|---|---|
| Cell gas | - | He | $H_2$ |
| $^{7}$Li [CPS per µg $L^{-1}$] | $14*10^3$ | - | - |
| $^{59}$Co [CPS per µg $L^{-1}$] | - | $4.5*10^3$ | $2.6*10^3$ |
| $^{89}$Y [CPS per µg $L^{-1}$] | $17*10^3$ | $3.4*10^3$ | $15*10^3$ |
| $^{205}$Tl [CPS per µg $L^{-1}$] | $13*10^3$ | $7.9*10^4$ | $12*10^4$ |
| average RSD [%] | 2.3 | 2.9 | 2.7 |
| $^{140}Ce^{16}O/^{140}Ce$ [%] | 1.3 | 0.5 | 1.3 |
| $^{140}Ce^{2+}/^{140}Ce^{+}$ [%] | 1.3 | 2.9 | 1.1 |

Table S2. Selected mass, tune mode, internal standard and detection limits (LoD)

| **Monitored isotope** | **Tune mode** | **Internal standard** | **Detection limit*** **(µg $L^{-1}$)** |
|---|---|---|---|
| $^{7}$Li | No-gas | $^{9}$Be | 0.025 |
| $^{11}$B | No-gas | $^{9}$Be | 0.025 |
| $^{23}$Na | He | $^{9}$Be | 27 |
| $^{24}$Mg | He | $^{9}$Be | 11 |
| $^{27}$Al | No-gas | $^{9}$Be | 2.4 |
| $^{31}$P | He | $^{9}$Be | 177 |
| $^{32}$S | He | $^{9}$Be | 302 |
| $^{39}$K | He | $^{9}$Be | 9.8 |
| $^{43}$Ca | He | $^{9}$Be | 257 |
| $^{51}$V | He | $^{74}$Ge | 0.02 |
| $^{53}$Cr | He | $^{74}$Ge | 0.08 |
| $^{55}$Mn | He | $^{74}$Ge | 0.55 |
| $^{56}$Fe | He | $^{74}$Ge | 1.4 |
| $^{59}$Co | He | $^{74}$Ge | 0.04 |
| $^{60}$Ni | He | $^{74}$Ge | 0.9 |

| | | | |
|---|---|---|---|
| $^{65}Cu$ | He | $^{74}Ge$ | 0.24 |
| $^{66}Zn$ | He | $^{74}Ge$ | 18 |
| $^{75}As$ | He | $^{74}Ge$ | 0.0030 |
| $^{78}Se$ | $H_2$ | $^{74}Ge$ | 0.01 |
| $^{85}Rb$ | He | $^{74}Ge$ | 0.023 |
| $^{88}Sr$ | He | $^{74}Ge$ | 0.1 |
| $^{98}Mo$ | No-gas | $^{74}Ge$ | 0.02 |
| $^{107}Ag$ | No-gas | $^{115}In$ | 0.001 |
| $^{111}Cd$ | No-gas | $^{115}In$ | 0.009 |
| $^{118}Sn$ | No-gas | $^{115}In$ | 0.02 |
| $^{121}Sb$ | No-gas | $^{115}In$ | 0.01 |
| $^{133}Cs$ | No-gas | $^{115}In$ | 0.002 |
| $^{137}Ba$ | No-gas | $^{115}In$ | 0.08 |
| $^{205}Tl$ | No-gas | $^{175}Lu$ | 0.0012 |
| $^{208}Pb$ | No-gas | $^{175}Lu$ | 0.03 |
| $^{238}U$ | No-gas | $^{175}Lu$ | 0.0003 |

$^{*}$LoD = $mean_{blanks} + 3*\sigma_{blanks}$

Table S3. Certified and determined values for elements in NIST SRM 1640a Trace Elements in Natural Water

| **Element** | **Cert. mass conc. [µg L$^{-1}$]** | | | **Analyzed mass conc. [µg L$^{-1}$] (n=4)** | | |
|---|---|---|---|---|---|---|
| Li | 0.4034 | ± | 0.0094 | 0.414 | ± | 0.029 |
| B | 300.7 | ± | 3.1 | 276.6 | ± | 4.6 |
| Na | 3137 | ± | 31 | 2985 | ± | 299 |
| Mg | 1058.6 | ± | 4.1 | 1018 | ± | 106 |
| Al | 53.0 | ± | 1.8 | 50.6 | ± | 3.7 |
| K | 579.9 | ± | 2.3 | 591 | ± | 78 |
| Ca | 5615 | ± | 21 | 5816 | ± | 725 |
| V | 15.05 | ± | 0.25 | 13.97 | ± | 0.82 |
| Cr | 40.54 | ± | 0.30 | 39.4 | ± | 1.7 |
| Mn | 40.39 | ± | 0.36 | 38.2 | ± | 1.7 |
| Fe | 36.8 | ± | 1.8 | 37.7 | ± | 3.2 |
| Co | 20.24 | ± | 0.24 | 18.85 | ± | 0.64 |
| Ni | 25.12 | ± | 0.14 | 23.27 | ± | 0.52 |
| Cu | 85.75 | ± | 0.51 | 85.20 | ± | 0.68 |
| Zn | 55.64 | ± | 0.35 | 52.1 | ± | 1.9 |
| As | 8.075 | ± | 0.070 | 7.79 | ± | 0.12 |
| Se | 20.13 | ± | 0.17 | 18.45 | ± | 0.97 |
| Rb | 1.198 | ± | 0.011 | 1.19 | ± | 0.14 |
| Sr | 126.03 | ± | 0.91 | 114.9 | ± | 8.5 |
| Mo | 45.60 | ± | 0.61 | 43.2 | ± | 2.1 |
| Ag | 8.081 | ± | 0.046 | 9.23 | ± | 0.30 |
| Cd | 3.992 | ± | 0.074 | 3.70 | ± | 0.18 |

| | | | | | | |
|---|---|---|---|---|---|---|
| Sb | 5.105 | ± | 0.046 | 4.733 | ± | 0.082 |
| Ba | 151.80 | ± | 0.83 | 140.0 | ± | 6.9 |
| Tl | 1.619 | ± | 0.016 | 1.439 | ± | 0.073 |
| Pb | 12.10 | ± | 0.05 | 11.20 | ± | 0.27 |
| U | 25.35 | ± | 0.27 | 24.21 | ± | 0.38 |

Table S4 Certified and determined values for elements in CRM 8414 Bovine Muscle Powder (*information values)

| **Element** | **Cert. mass conc. [mg kg$^{-1}$]** | | | **Analyzed mass conc. [mg kg$^{-1}$] (n=12)** | | |
|---|---|---|---|---|---|---|
| Na | 2100 | ± | 100 | 1799 | ± | 99 |
| Mg | 960 | ± | 95 | 842 | ± | 51 |
| Al* | 1.7 | ± | / | 1.14 | ± | 0.40 |
| P | 8360 | ± | 450 | 6911 | ± | 497 |
| S* | 8000 | ± | / | 6847 | ± | 266 |
| K | 15200 | ± | 400 | 13047 | ± | 884 |
| Ca | 145 | ± | 20 | 122 | ± | 10 |
| V* | 0.005 | ± | / | 0.00200 | ± | 0.00087 |
| Cr* | 0.071 | ± | / | 0.056 | ± | 0.011 |
| Mn | 0.37 | ± | 0.09 | 0.290 | ± | 0.021 |
| Fe | 71.2 | ± | 9.2 | 68.7 | ± | 2.8 |
| Co | 0.007 | ± | 0.003 | 0.00513 | ± | 0.00022 |
| Ni* | 0.05 | ± | / | 0.238 | ± | 0.017 |
| Cu | 2.84 | ± | 0.45 | 2.32 | ± | 0.39 |
| Zn | 142 | ± | 14 | 128.8 | ± | 5.0 |
| As | 0.009 | ± | 0.003 | 0.00732 | ± | 0.00044 |
| Se | 0.076 | ± | 0.010 | 0.0626 | ± | 0.0023 |
| Rb | 28.7 | ± | 3.5 | 24.7 | ± | 1.5 |
| Sr | 0.052 | ± | 0.015 | 0.0547 | ± | 0.0090 |
| Mo | 0.08 | ± | 0.06 | 0.0593 | ± | 0.0028 |
| Cd | 0.013 | ± | 0.011 | 0.0151 | ± | 0.0031 |
| Cs* | 0.05 | ± | / | 0.035 | ± | 0.013 |
| Ba* | 0.05 | ± | / | 0.0228 | ± | 0.0034 |
| Pb | 0.38 | ± | 0.024 | 0.32 | ± | 0.10 |

Table S5. Results of ANOVA and Kruskal-Wallis H test for 21 hives located in Mesić (n=336)

| Element | Independent t test | Kruskal-Wallis H test |
|---|---|---|
| B | $t(27)= 1.245, p=0.224$ | $\chi2(2)=0.441, p=0.507$ |

| | | |
|---|---|---|
| Na | t(27)= -6.131, p<0.0001 | χ2(2)=15.184, p<0.0001 |
| Mg | t(27)= -2.566, p=0.016 | χ2(2)=4.149, p=0.042 |
| Al | t(27)= 2.698, p=0.006 | χ2(2)=7.782, p=0.005 |
| P | t(27)= -5.473, p<0.0001 | χ2(2)=16.959, p<0.0001 |
| S | t(27)= -6.217, p<0.0001 | χ2(2)=19.608, p<0.0001 |
| K | t(27)= -2.494, p=0.019 | χ2(2)=5.931, p=0.015 |
| Ca | t(27)= 3.361, p=0.002 | χ2(2)=8.802, p=0.003 |
| V | t(27)= 3.922, p=0.001 | χ2(2)=13.508, p=0.0002 |
| Cr | t(27)= 1.254, p=0.221 | χ2(2)=2.684, p=0.101 |
| Mn | t(27)= 4.547, p=0.0001 | χ2(2)=16.596, p<0.0001 |
| Fe | t(27)= 3.468, p=0.002 | χ2(2)=7.782, p=0.005 |
| Co | t(27)= 3.910, p=0.001 | χ2(2)=9.608, p=0.002 |
| Ni | t(27)= 2.284, p=0.015 | χ2(2)=7.537, p=0.006 |
| Cu | t(27)= -2.269, p=0.031 | χ2(2)=4.331, p=0.037 |
| Zn | t(27)= -2.782, p=0.01 | χ2(2)=8.284, p=0.004 |
| As | t(27)= 3.572, p=0.001 | χ2(2)=9.067, p=0.003 |
| Se | t(27)= -12.490, p<0.0001 | χ2(2)=20.400, p<0.0001 |
| Rb | t(27)= 2.532, p=0.017 | χ2(2)=6.825, p=0.009 |
| Sr | t(27)= 4.555, p=0.0001 | χ2(2)=11.625, p=0.001 |
| Mo | t(27)= 6.970, p<0.0001 | χ2(2)=20.002, p<0.0001 |
| Ag | t(27)= 4.692, p<0.0001 | χ2(2)=15.882, p<0.0001 |
| Cd | t(27)= 4.805, p<0.0001 | χ2(2)=18.071, p<0.0001 |
| Sn | t(27)= 0.363, p=0.719 | χ2(2)=0.002, p=0.965 |
| Sb | t(27)= 3.808, p=0.001 | χ2(2)=12.549, p=0.0004 |
| Ba | t(27)= 4.787, p<0.0001 | χ2(2)=14.502, p=0.0001 |
| Pb | t(27)= 4.338, p=0.0001 | χ2(2)=12.549, p=0.0001 |
